# Abnormal microtubule dynamics disrupt nucleocytoplasmic transport in tau-mediated frontotemporal dementia

**DOI:** 10.1101/328187

**Authors:** F. Paonessa, L. Evans, R. Solanki, D. Larrieu, S. Wray, J. Hardy, S.P. Jackson, F.J. Livesey

## Abstract

The neuronal microtubule-associated protein tau (*MAPT*) is central to the pathogenesis of many dementias, including Alzheimer’s disease. Autosomal dominant mutations in *MAPT* cause inherited frontotemporal dementia (FTD), but the underlying pathogenic mechanisms are unclear. Using human stem cell models of FTD due to *MAPT* mutations, we find that tau becomes hyperphosphorylated and mislocalises to neuronal cell bodies and dendrites in cortical neurons, recapitulating a key early event in FTD. Mislocalised tau in the cell body leads to abnormal microtubule dynamics in FTD-MAPT neurons that grossly deform the nuclear membrane, resulting in defective nucleocytoplasmic transport. Neurons in the post-mortem human FTD-MAPT cortex have a high incidence of nuclear deformation, indicating that tau-mediated nuclear membrane dysfunction is an important pathogenic process in FTD. Defects in nucleocytoplasmic transport in FTD point to important commonalities in the pathogenic mechanisms of both tau-mediated dementias and ALS-FTD due to TDP-43 and C9orf72 mutations.

## INTRODUCTION

The microtubule-associated protein tau (MAPT, tau) is central to the pathogenesis of several different forms of dementia, including Alzheimer’s disease (AD), progressive supranuclear palsy, Pick’s disease, corticobasal degeneration and frontotemporal dementia (FTD) (Lee et al., 2001; Spillantini and Goedert, 2013). Frontotemporal dementia is the third most common cause of dementia, after Alzheimer’s disease and vascular dementia (Rossor et al., 2010). Autosomal dominant missense and splicing mutations in *MAPT* are causes of inherited or familial frontotemporal dementia (FTD-MAPT) (D’Souza et al., 1999; Goedert et al., 2012; Hutton et al., 1998). However, while it is well established that these mutations lead to hyperphosphorylation and aggregation of tau protein *in vivo* (Ballatore et al., 2007; Goedert et al., 2012), the cell biology of neuronal dysfunction and progressive neurodegeneration in this condition are currently not fully understood.

In healthy neurons, tau protein is almost exclusively localised to the axon, and several mechanisms have been suggested for its highly polarised cellular localisation, including selective mRNA and protein transport, local translation and local degradation (Y. Wang and Mandelkow, 2015). Mislocalisation and aggregation of tau in neuronal cell bodies are common features of tau-mediated dementias, including FTD and AD (Fu et al., 2016; Thies and Mandelkow, 2007; Zempel and Mandelkow, 2015). Protein aggregation is widely considered as inherently pathogenic in neurodegeneration (Fitzpatrick et al., 2017; Hernández-Vega et al., 2017), altering many cellular functions, most notably autophagy and proteostasis (Bence et al., 2001; Caballero et al., 2018; Lim and Yue, 2015). However, how *MAPT* mutations lead to tau hyperphosphorylation and mislocalisation, the effects of this mislocalisation on neuronal cell biology, and how this contributes to neuronal dysfunction and neurodegeneration, all remain poorly understood.

As a typical microtubule-binding protein, tau has several roles in regulating microtubule function and intracellular transport (Y. Wang and Mandelkow, 2015). Tau binds both alpha and beta tubulin subunits of microtubules, and has been demonstrated to both stabilise and promote microtubule growth (Kadavath et al., 2015; Witman et al., 1976). The presence of tau on microtubules can alter directions and rates of axonal transport (Dixit et al., 2008; Trinczek et al., 1999). Tau is a natively disordered protein and has recently been found to undergo fluid phase transitions at higher concentrations, nucleating microtubules when it does so (Hernández-Vega et al., 2017). Therefore, it is likely that the changes in tau levels, post-translational modifications and cellular localisation that occur in dementia lead to alterations in microtubule biology, particularly in the neuronal cell body.

To address the question of how *MAPT* mutations lead to neuronal dysfunction and neurodegeneration, we investigated the effects of two different classes of *MAPT* mutations on the cell biology of human iPSC-derived cortical neurons. We find that both missense and splicing *MAPT* mutations cause mislocalisation of tau to the cell bodies of neurons and marked changes in microtubule dynamics. Microtubules in the cell bodies of FTD-MAPT neurons actively deform the nuclear membrane, disrupting nucleocytoplasmic transport. Deficits in nuclear envelope function, including nucleocytoplasmic transport, are an important pathological process in ALS-FTD due to repeat expansions in *C9orf72* and *TDP-43* mutations (Chou et al., 2018; K. Zhang et al., 2018; 2015; Y.-J. Zhang et al., 2016). Our findings demonstrate that dysfunction of the nuclear membrane due to altered microtubule dynamics is a pathogenic process in dementias involving tau, expanding the group of neurodegenerative diseases that involve disrupted nucleocytoplasmic transport and suggesting common mechanisms of neuronal dysfunctional in these heterogeneous conditions.

## RESULTS

### Increased phosphorylation and altered cellular localisation of tau in FTD-MAPT neurons

To study the effects on neuronal cell biology of FTD-MAPT mutations, we generated excitatory cortical neurons (Shi et al., 2012b) from induced pluripotent stem cells (iPSC) derived from individuals with different autosomal dominant mutations in *MAPT* that are causal for early-onset FTD (Figs. 1 **& S1)**. We studied two different types of mutations: the *MAPT* IVS10+16 autosomal dominant mutation, which increases inclusion of exon 10, encoding the second microtubule-binding repeat, and thus altering the ratio of three (3R) and four (4R) tau isoforms (Hutton et al., 1998; Sposito et al., 2015); and the autosomal dominant *MAPT* P301L missense mutation that produces an aggregation-prone form of tau (Y. Wang and Mandelkow, 2015) **(Fig. S1)**.

**Figure 1.**
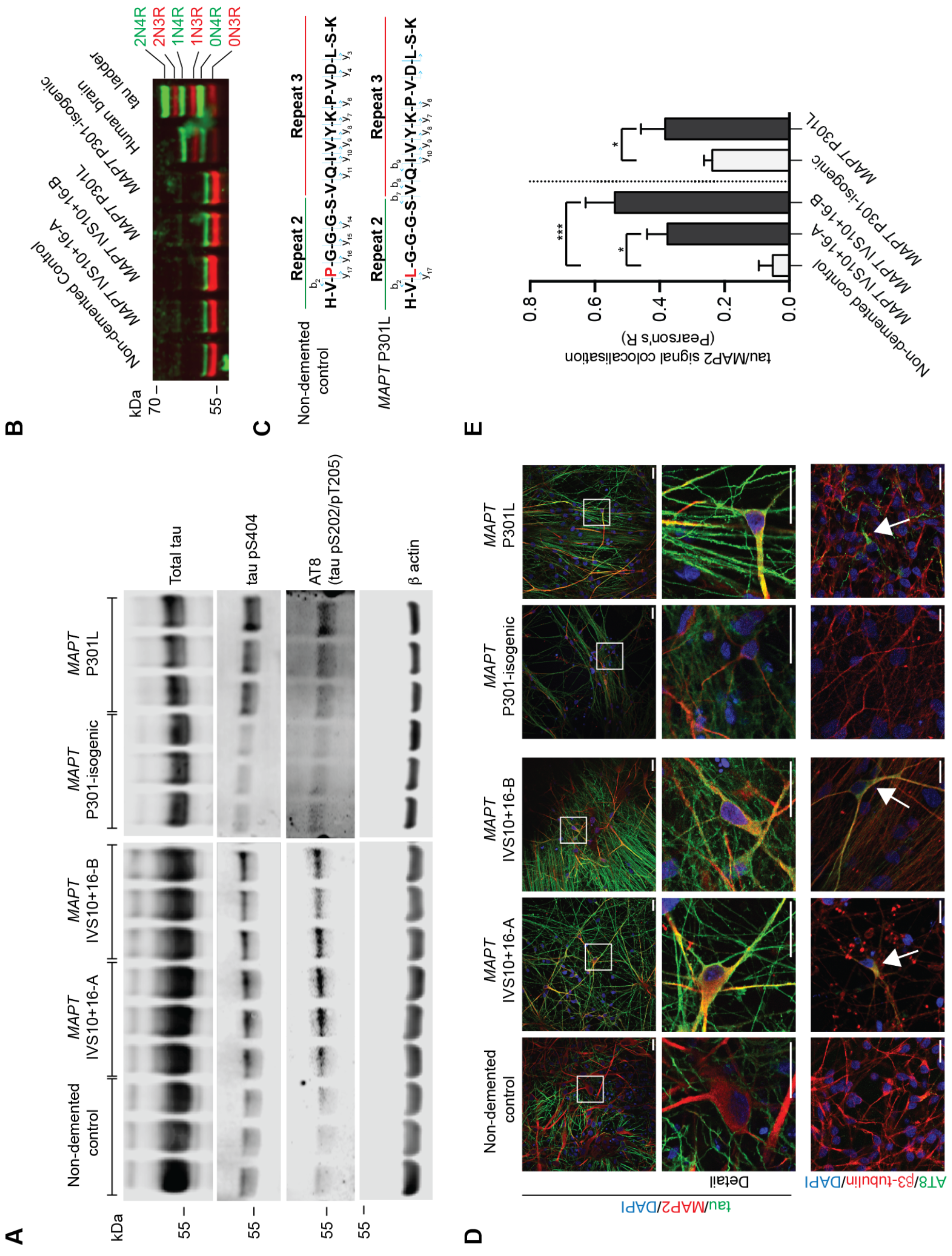
Increased phosphorylation and altered cellular localisation of tau in FTD-MAPT neurons. **(A)** Phosphorylated tau (pS404; AT8 [pS202/pT205]) is increased as a fraction of total tau (epitope 243-441) in FTD-MAPT neurons (*MAPT* IVS10+16-A/B and *MAPT* P301L) compared to non-demented and MAPT P301 isogenic control neurons (iPSC-derived neurons at 120 DIV; three biological replicates). β-actin was used as a loading control. Molecular weight (kDa) indicated. **(B)** Tau isoforms with three (3R; red) or four (4R; inclusion of region 2; green) microtubule binding regions were detected by western blot analysis of dephosphorylated protein extracts from iPSC-derived control and FTD-MAPT cortical neurons (120 DIV) and from post-mortem human cerebral cortex (non-demented individual). Tau isoforms were identified relative to a commercial tau ladder (Sigma). Molecular weight (kDa) indicated. **(C)** Peptide sequences identified by tau IP/mass spectrometry from iPSC-derived cortical neurons, confirming the inclusion of repeat 2 (corresponding to exon 10) of 4R tau. In *MAPT* P301L neurons, both proline and leucine were identified at position 301 (highlighted red). See also Figure S2. **(D)** Tau protein is mislocalised to MAP2-positive cell bodies and dendrites in iPSC-derived FTD-MAPT neurons. Confocal images of iPSC-derived control and FTD-MAPT neurons, (120 DIV; tau, green; MAP2, red; DAPI, blue). Hyperphosphorylated, AT8-positive tau (AT8; green) is found in cell bodies of FTD-MAPT neurons (arrows), but not in controls (β3-tubulin, red; DAPI, blue). Scale bars = 20 μm. **(E)** Increased co-localisation of tau and MAP2 protein in FTD-MAPT neurons, compared to non-demented control neurons, analysed by Pearson’s R correlation (control lines: grey bars; FTD-MAPT lines: black; significance was determined for three sample comparison of non-demented control and two *MAPT* IVS10+16 lines using one-way ANOVA followed by Tukey’s test; * = P<0.05, *** = P<0.001. Pair-wise comparison of the *MAPT* P301L line and its isogenic control was carried out using Student’s *t* test; *=P<0.05; error bar represents s.e.m.; n = 3 independent experiments). See also Figure S1.

Total tau content was similar in neurons of each genotype, collectively referred to here as FTD-MAPT neurons **(Fig. 1A)**. However, tau phosphorylation was increased in FTD-MAPT neurons compared to controls **(Fig. 1A)**, notably at Ser404 and Ser202/Thr205 (AT8), epitopes typically hyper-phosphorylated in tau-mediated dementias (Alonso et al., 2004; J.-Z. Wang et al., 2013). As both mutations are dependent on expression of exon 10 of *MAPT*, we confirmed translation of exon 10 in neurons generated from all iPSC lines by western blotting and mass spectrometry (Fig. 1B, C **& S1)**.

Mislocalisation of tau from axons to neuronal cell bodies and dendrites is an early event in FTD *in vivo* (Götz et al., 1995; Hoover et al., 2010; Kowall and Kosik, 1987). As expected, control neurons showed a predominantly axonal distribution of tau, with tau largely absent from MAP2-positive neuronal cell bodies and dendrites **(Fig. 1D)**. In contrast, tau was commonly present in MAP2-positive cell bodies and dendrites in both *MAPT* IVS10+16 and P301L neurons **(Fig. 1D, E)**. Furthermore, tau within cell bodies and dendrites of FTD-MAPT neurons was hyperphosphorylated, as detected by AT8-immunoreactivity (phospho-S202/T205; **Fig. 1D)**.

### Microtubules invade the nucleus in FTD-MAPT neurons

Given the mislocalisation of tau to the cell bodies of FTD-MAPT neurons, we studied neuronal microtubule dynamics in control and FTD-MAPT neurons (Figs. 2 **& S2)**. Actively extending microtubules were live-imaged in iPSC-derived neurons of each genotype by expression of GFP-tagged EB3 **(Fig. 2**; **MovieS1)**, the microtubule plus-end binding protein (+TIP; (Akhmanova and Steinmetz, 2008)). Average microtubule dynamics were not different between non-demented control and FTD-MAPT neurons, with similar rates of extensions and retractions measured among the various genotypes **(Table S1)**.

**Figure 2.**
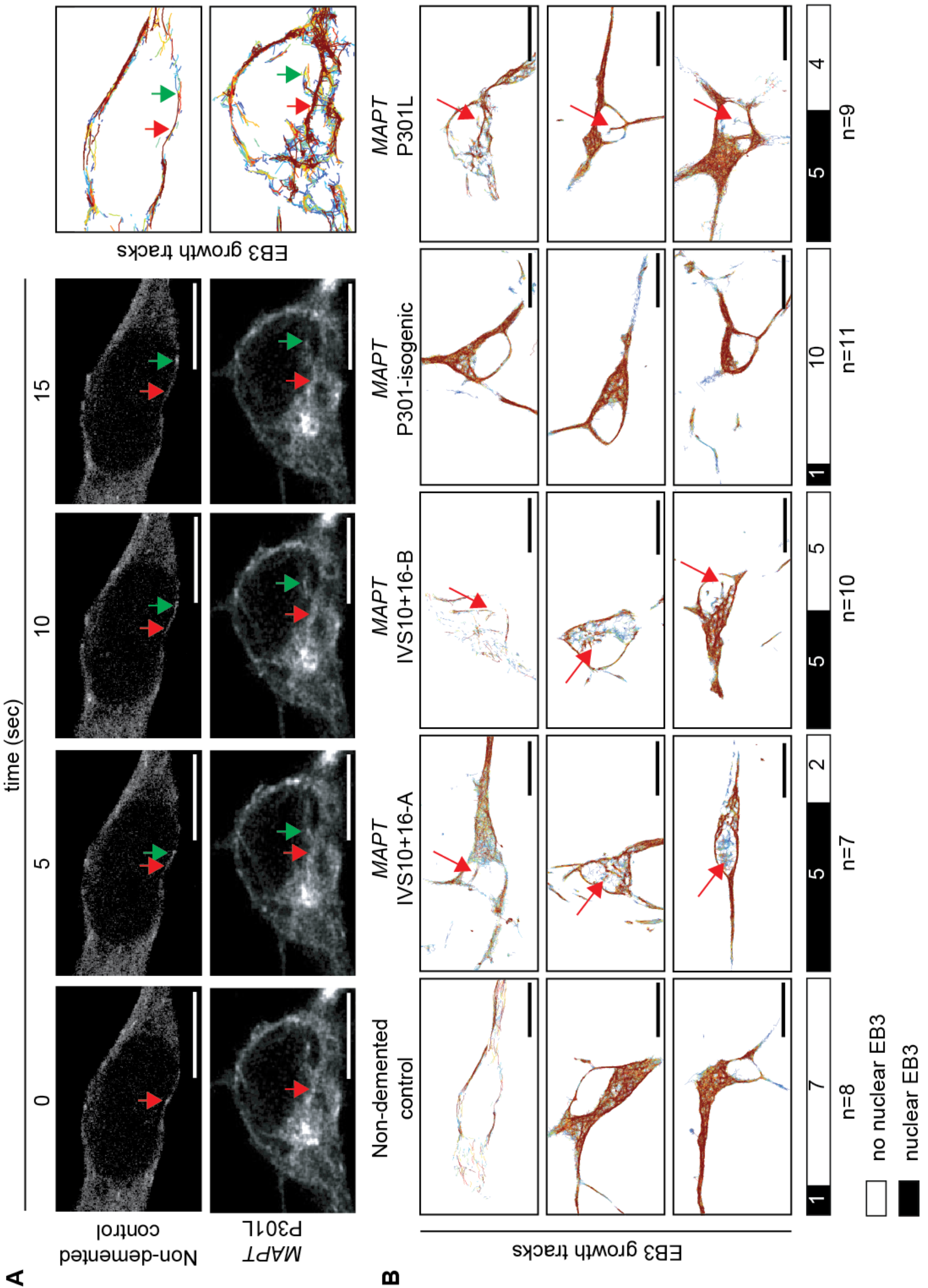
Microtubules invade the nucleus in FTD-MAPT neurons. **(A)** Representative stills (5-second intervals) from GFP-EB3 live imaging of nondemented iPSC-derived control and *MAPT* P301L neurons (120 DIV) show actively growing microtubules extending into the nucleus in the *MAPT* mutant neurons. Red arrows indicate the position of an EB3-GFP comet at time point 0 (far left); green arrows indicate the comet position in each subsequent frame. Right panels show cumulative tracks (growth tracks) of EB3 comets over the 200-second interval. **(B)** Total microtubule trajectories (200-second interval) demonstrate multiple microtubule extensions into the nuclei of FTD-MAPT neurons (*MAPT* IVS10+16-A/B and *MAPT* P301L), compared to non-demented and MAPT P301 isogenic control neurons. Red arrows indicate examples of trajectories within the nucleus. Bars indicate the number of sampled neurons with (black), and without (white), nuclear EB3 growth tracks; n=number of imaged neurons. Scale bars = 10μm. See also Video S1 and Table S1. See also Figure S2.

However, microtubule dynamics were qualitatively different in the cell bodies of control neurons compared with FTD-MAPT neurons. Non-demented control neurons typically had many actively growing microtubules within the cell body that extended around a smooth, oval nucleus **(Fig. 2A, B**; **Movie S1)**. In contrast, many FTD-MAPT neurons had microtubules with plus ends projecting into the nucleus (15/26 FTD-MAPT neurons; Fig. 2), an event that was infrequently detected in both groups of control neurons (2/19; Fig. 2). Notably, those microtubules that abnormally projected into the nucleus in FTD-MAPT neurons frequently originated from a pronounced focus that resembled a microtubule organising centre **(Fig. 2A, B**; **Movie S1)**.

### Microtubules deform the nuclear envelope in FTD-MAPT neurons

Microtubules couple to the nuclear membrane through the LINC complex (Crisp et al., 2006). This physical association results in transmission of mechanical forces that influence nuclear shape and integrity (Chang et al., 2015), affecting the function of the nuclear envelope (Webster et al., 2009). Given the abnormal projection of microtubules into the nucleus in FTD-MAPT neurons, we studied the shape of the nuclear envelope in iPSC-derived neurons. Marked differences were present in nuclear shape between non-demented controls and FTD-MAPT neurons, as demonstrated by large folds, or invaginations, of the laminB1-positive inner nuclear lamina within the nucleus (**Fig. 3**). Quantification of the proportions of neurons with deformation of the nuclear membrane, as defined by the presence of LaminB1-positive domains within the nucleus (**Fig. 3B;** see Fig S3, Methods and Supplementary Materials for details of quantification), found that approximately 25% of *MAPT* P301L and 40% of *MAPT* IVS10+16 neurons have deep nuclear invaginations, compared with less than 10% of control neurons **(Fig. 3A, B)**.

**Figure 3.**
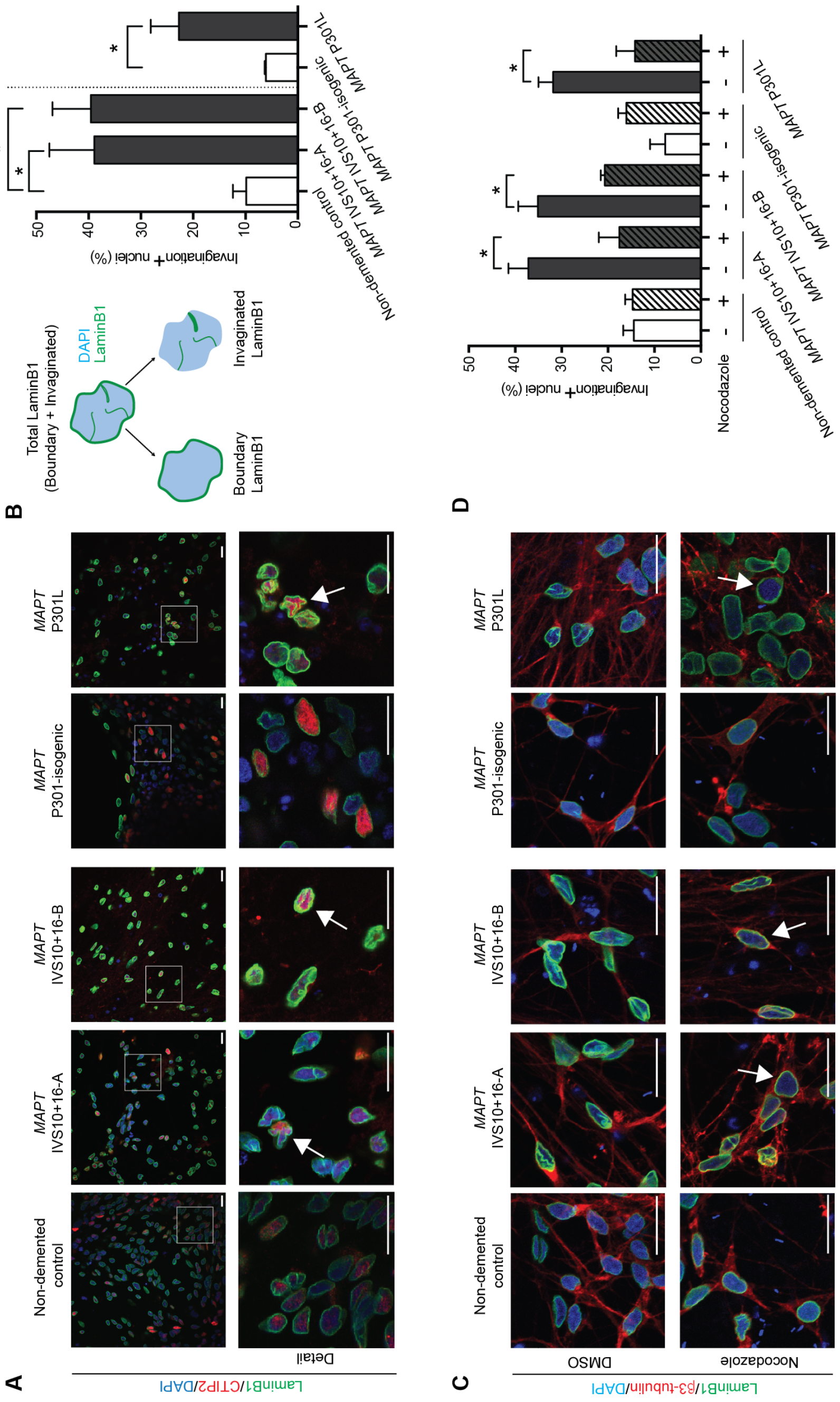
Microtubules deform the nuclear envelope in FTD-MAPT neurons. **(A)** Marked abnormalities of nuclear lamina shape in FTD-MAPT neurons. Confocal images of the nuclear lamina (laminB1, green) in FTD-MAPT neurons (*MAPT* IVS10+16-A/B and *MAPT* P301L; neuronal transcription factor CTIP2, red) compared to non-demented and MAPT P301 isogenic control neurons (120 DIV). White arrows indicate examples of nuclei with pronounced deformation of the nuclear lamina. **(B)** FTD-MAPT neurons have increased numbers of cells with deformed nuclear membranes, as defined by the shape of the inner nuclear lamina. Cartoon of image analysis method used to quantify nuclear invaginations, as a measure of distortion of the nuclear membrane: nuclear area was established using DAPI (blue), and nuclear lamina (LaminB1; green) was assigned as either nuclear boundary or invaginated (i.e., within the nucleus). The fraction of total laminB1 that was invaginated was used empirically to define a threshold for defining neurons as having nuclear membrane invaginations (see Figure S3 for details). Between 25% (*MAPT* P301L) and 40% (*MAPT* IVS10+16) of FTD-MAPT neurons have nuclear invaginations, compared with less than 10% of control neurons. Significance was determined for three sample comparison of non-demented control and two *MAPT* IVS10+16 lines using one-way ANOVA followed by Tukey’s test; * = P<0.05; pair-wise comparison of the *MAPT* P301L line and its isogenic control was carried out using Student’s *t* test; *=P<0.05; error bar represents s.e.m.; n = 3 independent experiments. **(C)** Acute depolymerisation of microtubules reverses nuclear lamina invaginations and restores rounded nuclear shapes. Confocal images of control and FTD-MAPT neurons (using genotypes described in A; 120 DIV), treated with DMSO (vehicle) or 10 μM nocodazole for 3 hours (laminB1, green; β3-tubulin, red; DAPI, blue). Scale bars = 10 μm. **(D)** The proportion of FTD-MAPT neurons with nuclear lamina invaginations is significantly reduced by nocodazole treatment. Quantification of neurons with abnormalities of the nuclear lamina was carried out as in B. n = 3 independent experiments; Error bars = s.e.m. Significance was determined using one-way ANOVA followed by Tukey’s test; * = P<0.05; error bar represents s.e.m.; n = 3 independent experiments). See also Figure S3.

Confirming that microtubules actively deformed the nucleus in FTD-MAPT neurons, we found that acute microtubule depolymerisation (with the small molecule nocodazole) reduced the number of nuclear invaginations and restored round nuclear morphology **(Fig. 3C, D)**. We conclude that the pronounced deformations of the neuronal nuclear membrane in FTD-MAPT neurons are actively mediated by microtubules.

### Super-resolution imaging demonstrates close apposition of tau and tubulin within nuclear lamina invaginations

To further study the spatial relationships between tau, microtubules and the nuclear envelope, we conducted a detailed analysis of the neuronal nucleus in iPSC-derived neurons using 3D-STED super-resolution imaging. 3D-STED imaging demonstrated that invaginations of the nuclear lamina present in FTD-MAPT neurons commonly extended deeply into the nucleus, in some cases traversing the entire length of the nucleus, forming pronounced folds **(Fig. 4A)**. In comparison, nuclei from non-demented controls had a regular, smooth morphology, with few examples of nuclear lamina invaginations **(Fig. 4A)**.

**Figure 4.**
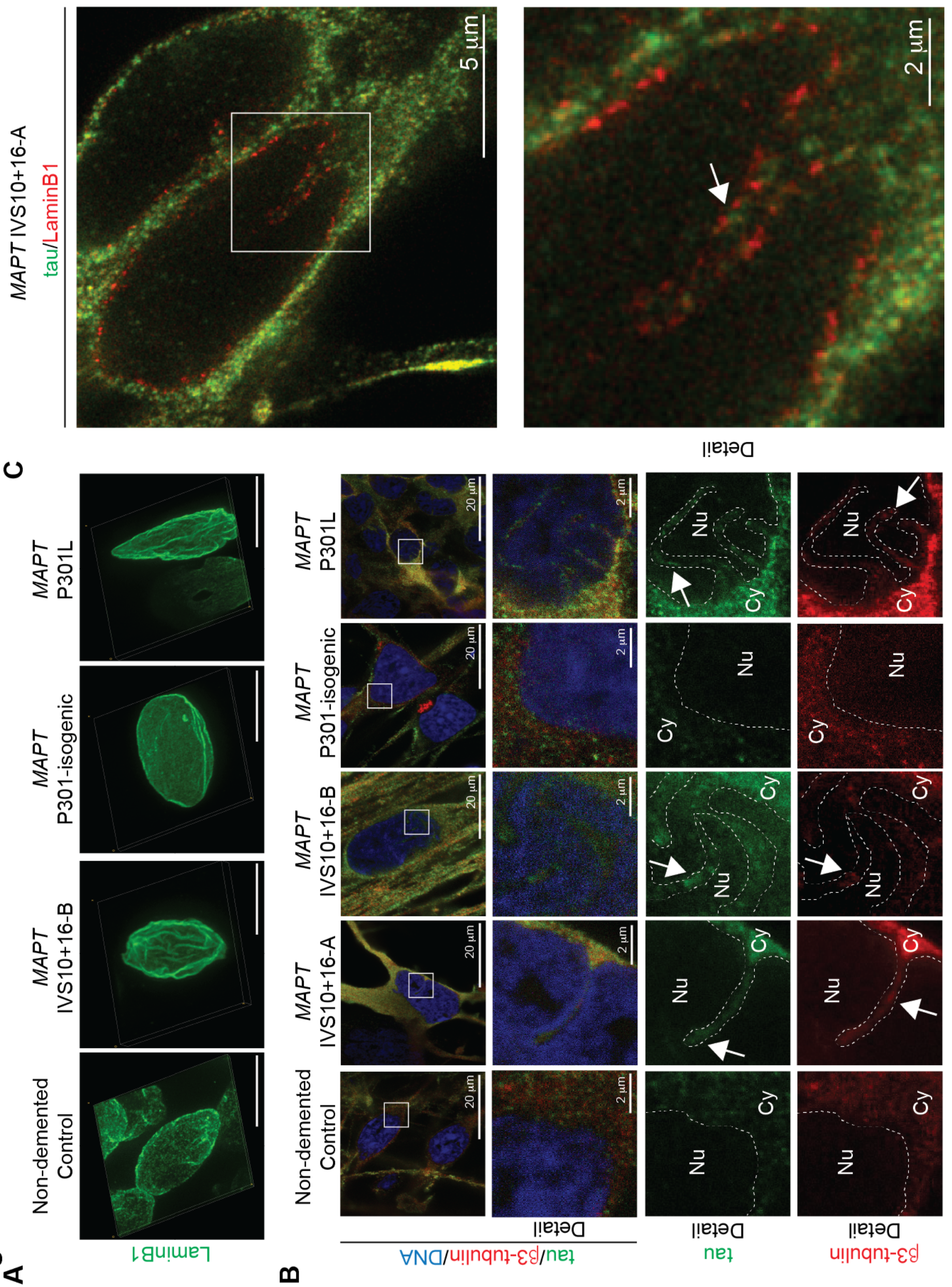
Super-resolution imaging demonstrates close apposition of tau and tubulin within nuclear lamina invaginations. **(A)** 3D-reconstructions using STED imaging of the nuclear lamina (laminB1; green) in FTD-MAPT iPSC-derived neurons (*MAPT* IVS10+16 and *MAPT* P301L) reveal pronounced nuclear invaginations compared with non-demented and MAPT P301-isogenic control neurons (120 DIV). Scale bar = 10μm. **(B)** Tau is in close proximity to the nuclear lamina within nuclear invaginations of FTD-MAPT neurons. STED imaging of control and FTD-MAPT iPSC-derived neurons (using genotypes described in A; 120 DIV; tau, green; β3-tubulin, red; DNA, Yo-Pro, blue). Detail from white boxes in upper panels, showing both merge of all channels, and single channel images of tau (green) or β3-tubulin (red). Dashed lines indicate the boundary between the nucleus (Nu) and cytoplasm (Cy). Arrows indicate invaginations into the nucleus. **(C)** Nuclear invaginations are lined with nuclear lamina and contain tau. STED imaging of *MAPT* IVS10+16-A neurons (120DIV; tau, green; and laminB1, red). Arrow indicates tau within a nuclear invagination, in close proximity to the laminB1-positive inner nuclear lamina.

As observed by confocal microscopy, tau protein was found to be abundant in the cell bodies of FTD-MAPT neurons by STED imaging, and in those neurons was closely apposed to the outer nuclear membrane **(Fig. 4B)**. Both tau protein and neuronal tubulin were found within nuclear lamina invaginations in FTD-MAPT neurons **(Fig. 4B)**. STED imaging demonstrated that tau within nuclear membrane invaginations is within 100s of nanometres of the nuclear lamina **(Fig. 4C)**. Given that laminB1 filaments line the inner surface of the nuclear envelope, which is typically of the order of 15-60nm in width (Burke and Stewart, 2013; Gerace and Huber, 2012), we conclude that tau is in close proximity to proteins in the outer membrane of the nuclear envelope in FTD-MAPT neurons.

### Tau-containing nuclear lamina invaginations in neurons of the post-mortem FTD-MAPT cerebral cortex

Having identified nuclear lamina defects in iPSC-derived FTD-MAPT neurons, we asked whether alterations of the nuclear lamina are also a feature of FTD-MAPT *in vivo*. To do so, we studied the incidence of invaginations of the nuclear lamina in the frontal and temporal cortex of two individuals diagnosed with FTD due to *MAPT* IVS10+16 mutations. We quantified the fraction of all nuclei with nuclear invaginations, in both neuronal and non-neuronal cells, within each brain region. Invaginations of the nuclear lamina, with intranuclear inclusions of laminB1, were found in both non-demented control individuals, but were more frequent in cells of the deep cortical layers in *postmortem* cerebral cortex from individuals with FTD due to the *MAPT* IVS10+16 mutation (**Fig. 5A, B**; **Fig. S4)**. This was the case in both frontal and temporal cortex **(Fig. 5B)**.

**Figure 5.**
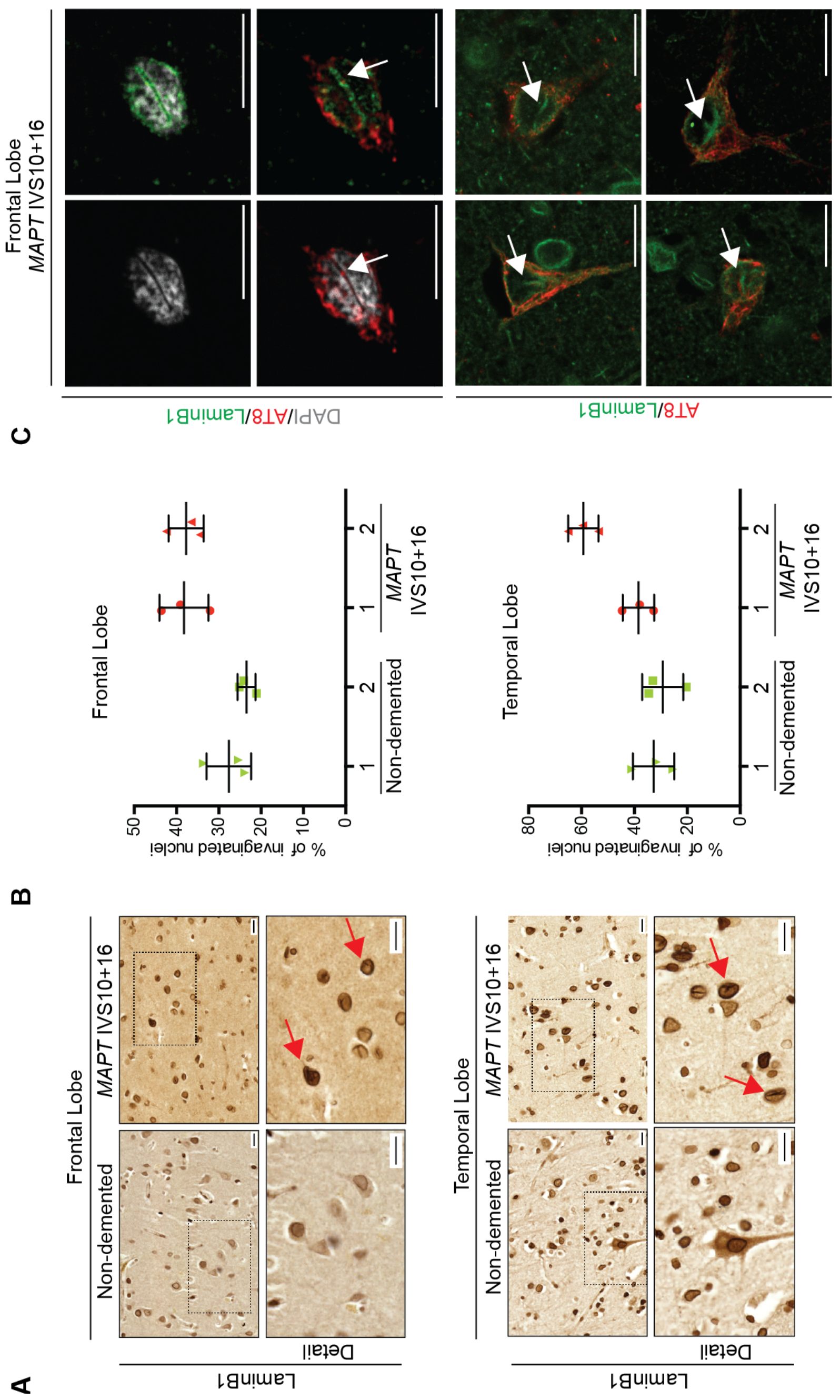
Tau-containing nuclear lamina invaginations in neurons of the postmortem FTD-MAPT cerebral cortex. **(A)** Increased incidence of laminB1-positive nuclear invaginations *in vivo*, in postmortem FTD-MAPT IVS10+16 cortex compared to age-matched controls. Immunohistochemistry of laminB1 in (upper panels) frontal and (lower panels) temporal cortex from individuals with FTD due to the *MAPT* IVS10+16 mutation or age-matched controls (non-demented). Arrows indicate nuclei exhibiting nuclear invaginations. **(B)** Percentage of invaginated nuclei in deep layers of frontal and temporal cortex of two control (green) and two *MAPT* IVS10+16 individuals (red), calculated from 20 individual imaging fields, Nuclei were scored by three observers, blinded to the identity of the post-mortem samples, and the averages of the three measurements are shown. Error represents s.e.m. **(C)** Hyperphosphorylated, AT8+ tau within nuclear lamina invaginations in neurons of the frontal cortex of individuals carrying *MAPT* IVS10+16 mutation. Representative neuron (upper panels), showing an extensive nuclear invagination (laiminB1, green; DAPI, grey) containing hyperphosphorylated tau (AT8, red). Further examples of frontal cortex neurons with nuclear invaginations containing phosphorylated tau; laminB1 (green) and phosphorylated tau (AT8; red). Arrows indicate nuclear invaginations. Scale bars = 10 μm. See also Figure S4.

Our data from iPSC-derived FTD-MAPT neurons predicted that laminB1-positive nuclear invaginations would be associated with the presence of phosphorylated tau within the neuronal cell body. Consistent with this, we found that the nuclear lamina was grossly disrupted in neurons that had high levels of hyperphosphorylated (AT8+) tau and neurofibrillary tangles in the *post-mortem* FTD-MAPT IVS10+16 cerebral cortex **(Fig. 5C, F**; **Fig. S4)**, and those neurons frequently contained pronounced nuclear lamina invaginations **(Fig. 5C)**. Furthermore, nuclear lamina invaginations in such neurons also commonly contained AT8-positive, hyperphosphorylated tau **(Fig. 5C)**.

### Disrupted nucleocytoplasmic transport in FTD-MAPT neurons

Abnormalities of the nuclear lamina such as those reported here are also found in ageing diseases, such as Hutchinson-Gilford progeria syndrome (Broers et al., 2006). Nuclear membrane distortion in response to mechanical forces leads to deleterious effects on many aspects of nuclear function, disrupting nucleocytoplasmic transport (Kelley et al., 2011). We confirmed that the nuclear lamina/membrane invaginations present in iPSC-derived FTD-MAPT neurons also contained nuclear pores within these membrane folds, with nuclear pores (labelled by NUP98) co-localising with laminB1-positive invaginations **(Fig. 6A)**.

**Figure 6.**
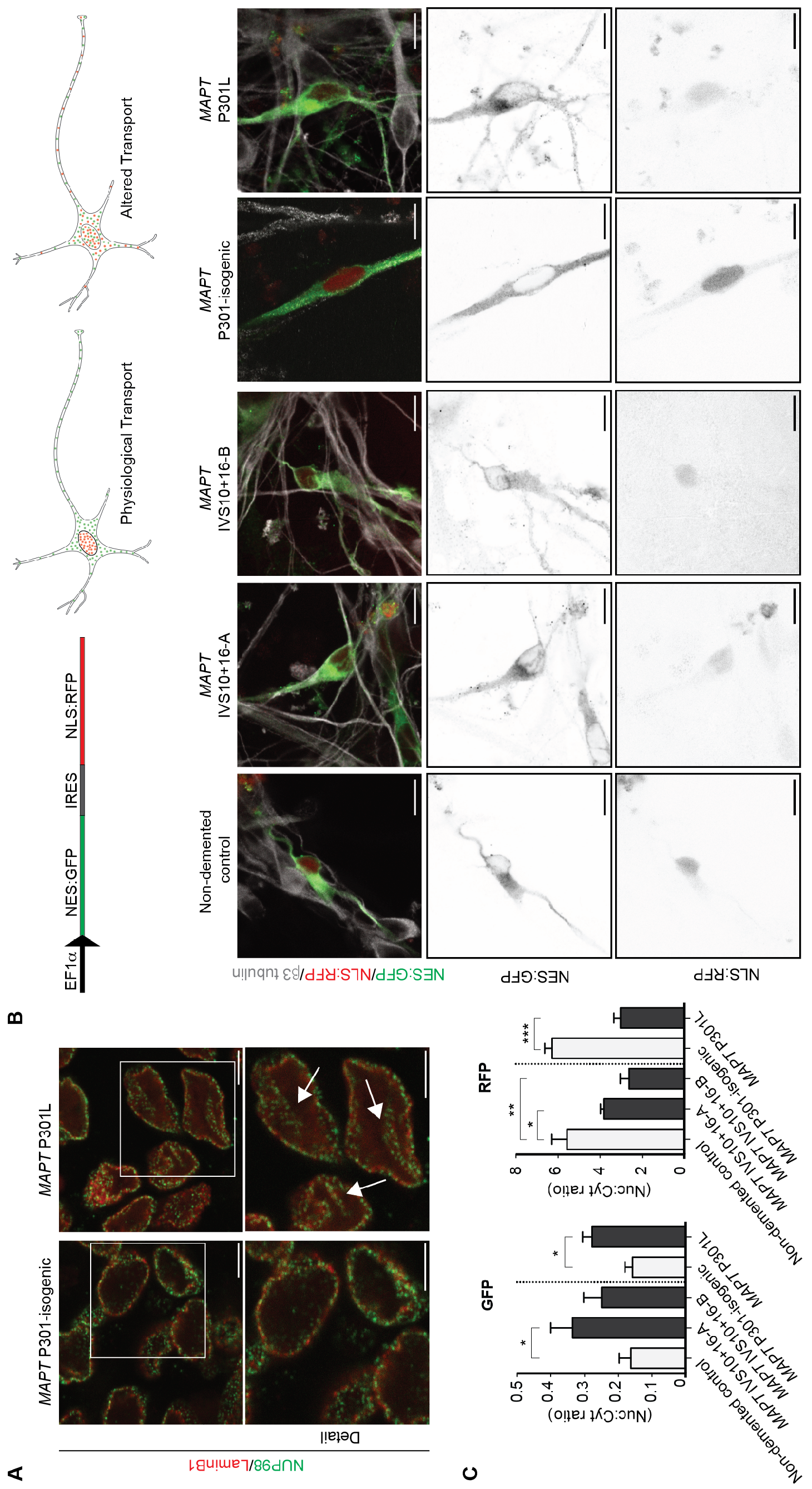
Disrupted nucleocytoplasmic transport in FTD-MAPT neurons. **(A)** Nuclear pores are clustered within nuclear membrane invaginations in FTD-MAPT neurons. Nuclear pore subunit NUP98 (green) remains co-localised with laminB1 (red) within nuclear invaginations in *MAPT* P301-isogenic neurons and *MAPT* P301L, (120 DIV). Arrows indicate nuclear membrane invaginations. Scale bars = 5 μm. **(B)** Functional assay demonstrates disrupted nucleocytoplasmic transport in human iPSC-derived FTD-MAPT neurons. Schematic illustrates lentiviral vector for coexpression of NES:GFP (nuclear export signal fused to GFP) and NLS:RFP (nuclear localisation signal fused to RFP) within human neurons and relative distributions of both proteins in healthy neurons and in cells with defective nucleocytoplasmic transport. Representative confocal images of control and FTD-MAPT neurons, (*MAPT* IVS10+16-A/B and *MAPT* P301L; all 120 DIV) expressing GFP:NES and RFP:NLS (GFP, green; RFP, red; β3 tubulin). Greyscale images of NES-GFP and NLS-RFP localisation in representative cells of each genotype are shown: FTD-MAPT neurons show an increase of GFP within the nucleus, and a reduction in nuclear localisation of NLS-RFP. Scale bars = 20 μm. **(C)** Quantification of the nuclear:cytoplasmic ratio for both NES:GFP and NLS:RFP demonstrate altered nuclear transport in FTD-MAPT genotypes relative to controls: NES:GFP is increased in the nucleus of FTD-MAPT neurons, whereas NLS:RFP is decreased (Significance was determined for three sample comparison of non-demented control and two *MAPT* IVS10+16 lines using one-way ANOVA followed by Dunnett’s test; * = P<0.05; ** = P<0.01; *** = P<0.001. Pair-wise comparison of the *MAPT* P301L line and its isogenic control was carried out using Student’s *t* test; *=P<0.05; error bar represents s.e.m.; n = 4 independent experiments). See also Figure S5.

To assess whether alterations in the nuclear membrane in FTD-MAPT neurons result in defects in nucleocytoplasmic transport, we expressed NES:GFP and NLS:RFP from a single construct in iPSC-derived neurons (Mertens et al., 2015). This assay enables measurement of the integrity of both nuclear localisation/accumulation and cytoplasmic retention/nuclear exclusion within individual neurons (Fig. 6B **& S5)**. Control iPSC-derived neurons had discrete cellular distributions of each protein, with prominent nuclear RFP and cytosolic GFP **(Fig. 6B)**. In contrast, localisation of NLS:RFP was altered in FTD-MAPT neurons such that there was a marked decrease in the nuclear:cytoplasmic RFP ratio **(Fig. 6B, C)**. Conversely, nuclear exclusion of NES:GFP was reduced in FTD-MAPT neurons, with an increase of GFP within the nucleus **(Fig. 6B, C)**. Together, these data demonstrate defects in the selective permeability of the nuclear envelope in FTD-MAPT neurons, indicating a general failure of nucleocytoplasmic transport within FTD-MAPT neurons.

## DISCUSSION

The cell and molecular biology of the pathogenesis of frontotemporal dementia due to *MAPT* mutations are not well understood. Currently, it is thought that *MAPT* mutations all lead to tau protein aggregation, and that protein aggregation is the primary driver of neurodegeneration (Ballatore et al., 2007; Spillantini and Goedert, 2013). However, protein aggregation may only represent the late stage of the disease, and the processes preceding and leading to neurofibrillary tangle formation and cellular dysfunction remain to be elucidated. Here we report the use of human stem cell systems, combined with *MAPT* mutations, to study the effects of those mutations on neuronal cell biology, finding that tau-mediated dementias lead to defective neuronal nucleocytoplasmic transport.

Focusing on two different types of *MAPT* mutations causal for FTD, we have found that both IVS10+16 and P301L mutations lead to marked defects in nucleocytoplasmic transport in human neurons. We find that both missense and splicing mutations in *MAPT* alter tau protein localisation and phosphorylation within iPSC-derived neurons within four months in cell culture, recapitulating a well-described aspect of early FTD pathology *in vivo* (Götz et al., 1995; Hoover et al., 2010; Kowall and Kosik, 1987). Mislocalisation of tau in the cell bodies of FTD-MAPT neurons in culture leads to marked changes in microtubule dynamics, causing deformation of the nuclear membrane both in cell culture and in the human FTD-MAPT cortex *in vivo*. Disruption of the nuclear lamina is commonly associated with dysfunction of the nuclear envelope, and we find marked disruption of nucleocytoplasmic transport in FTD-MAPT neurons. Together, these data indicate that perturbation of the function of the nuclear membrane and disruption of nucleocytoplasmic transport is an important pathological process in FTD due to *MAPT* mutations.

An early event in FTD is the mislocalisation of tau from axons to cell bodies and dendrites, and this key stage in disease progression is also an early event in iPSC-derived models of FTD-MAPT. *In vivo*, mislocalisation of tau is typically associated with tau hyperphosphorylation (Götz et al., 1995; Spillantini and Goedert, 2013). We find this also occurs in iPSC-derived FTD-MAPT neurons, where we detected increased tau phosphorylation at serine 202/threonine 205 (the AT8 epitope) and also at serine 404. The sequence in which mislocalisation and hyperphosphorylation take place in FTD in vivo, and in iPSC-derived FTD-MAPT neurons in culture, is not currently clear, nor are the mechanisms by which these processes occur. The appearance of tau within cell bodies and dendrites indicates a breakdown of the cellular polarity mechanisms that maintain the axonal enrichment of tau protein and its exclusion from the somatodendritic compartment, mechanisms which are poorly understood.

The two heterozygous, dominant *MAPT* mutations studied here have different effects on tau protein in neurons. The *MAPT* P301L missense mutation, like many missense mutations in the microtubule binding region domain of *MAPT*, increases the tendency of tau to aggregate in cell-free assays and in transgenic mouse models (Bergen et al., 2005; Lewis et al., 2000; Shammas et al., 2015). In contrast, the IVS10+16 mutation is not a coding mutation, but rather is an intronic single base change that favours the inclusion of exon 10 in the *MAPT* mRNA, increasing the amount of tau containing four microtubule-binding repeats (4R), relative to the three-repeat (3R) form (Hutton et al., 1998). However, despite these differences, the changes in the forms of tau in both *MAPT* P301L and IVS10+16 neurons both lead to mislocalisation and increased phosphorylation of tau. This finding suggests that either the presence of a pool of P301L tau, or a shift in the 3R:4R tau ratio, alter a common pathway that regulates tau distribution within neurons, tau phosphorylation, or both.

In both *MAPT* IVS 10+16 and P301L mutant neurons, the appearance of tau in cell bodies is accompanied by marked qualitative changes in neuronal microtubule dynamics. Microtubules in FTD-MAPT neurons actively deform the nuclear envelope, which we find can be reversed by depolymerisation of microtubules. Tau has multiple roles in stabilising microtubules (Y. Wang and Mandelkow, 2015), and microtubules are coupled to the nuclear membrane through the LINC complex (Chang et al., 2015; Crisp et al., 2006; Luo et al., 2016). Therefore, it is likely that the overall effect of the presence of tau in the cell body is to promote microtubule stability, leading to increased pushing forces on the nuclear membrane and the formation of invaginations in the nuclear membrane. As tau has recently also been found to promote microtubule nucleation when undergoing phase transitions at high concentration (Hernández-Vega et al., 2017), accumulation of tau in the cell body may also lead to a net increase in microtubule growth events and increased pushing forces on the nucleus by facilitating microtubule nucleation.

Alterations in nuclear shape and nuclear membrane function are a common feature of cellular ageing, including in the nervous system, and are associated with multiple deleterious changes in nuclear biology, including chromatin changes and disrupted nucleocytoplasmic transport (Frost, 2016; Oberdoerffer and Sinclair, 2007). *Drosophila* models of FTD, with neuronal expression of human *MAPT* R406W, have nuclear shape abnormalities and chromatin changes (Frost et al., 2016). Perturbations of the nuclear lamina have been described in the *post-mortem* AD brain (Frost et al., 2016), including the juxtaposition of neurofibrillary tangles of tau with the nuclear membrane (Sheffield et al., 2006). We also find here an increase in nuclear invaginations in neurons of the human post-mortem *MAPT* IVS10+16 cortex, and the presence of hyperphosphorylated tau within nuclear invaginations in tangle-bearing neurons. Together, these different studies are consistent with a pathological effect of tau within the neuronal cell body in FTD and AD, where the presence of tau alters microtubule biology, resulting in pronounced abnormalities of the neuronal nucleus and defective nucleocytoplasmic transport.

Microtubule deformation of the nucleus is a phenotype also seen in classic laminopathies such as the accelerated aging disorder Hutchinson-Gilford Progeria Syndrome (HGPS), where the primary defect is due to mutant lamin A/C protein (Capell and Collins, 2006). In that case, microtubules also contribute to nuclear deformations, leading to defects in nucleocytoplasmic transport (Kelley et al., 2011; Larrieu et al., 2014; Snow et al., 2013). Changes in nuclear envelope function in other neurodegenerative diseases, including ALS-FTD due to repeat expansions in *C9orf72* and Huntington’s disease, have recently been reported (Freibaum et al., 2015; Grima et al., 2017; Jovičić et al., 2015; K. Zhang et al., 2015). Our finding here of disruption of the neuronal nuclear membrane as a consequence of *MAPT* mutations in frontotemporal dementia extends this pathogenic mechanism to dementias where protein aggregation has been thought to be the primary driver of neurodegeneration. These data suggest that dysfunction of the nuclear membrane may be a common pathogenic process in diverse neurodegenerative diseases, which could be targeted therapeutically with agents that regulate microtubule functions, nucleocytoplasmic transport and/or associated processes.

## METHODS

### Generation of iPSC-derived cortical neurons and drug treatments

*MAPT* IVS10+16-A and *MAPT* IVS10+16-B mutant iPSCs were as reported in (Sposito et al., 2015). *MAPT* P301L was generated from Janssen Pharmaceutica NV by TALEN editing the line *MAPT* P301-isogenic, under the IMI STEMBANCC project agreement ICD 483960. The non-demented control line was previously reported (Israel et al., 2012). Pluripotent cells were cultured as previously described (Beers et al., 2012). Differentiation of iPSCs to cortical neurons was carried out as described, with minor modifications (Shi et al., 2012b; 2012a). For nocodazole (Tocris) treatment, neurons were grown for 120 days *in vitro* (DIV) and compound was added at 33 μM for 4 hr before imaging.

### Protein extraction and western blot analysis

Total cell protein was extracted using RIPA buffer (Sigma; R0278) supplemented with protease inhibitors (Sigma; 4693159001) and Halt phosphatase inhibitors (Thermo Scientific; 78420). Immunoblots were detected using LI-COR Odyssey CLx Infrared Imaging System and processed with the Image Studio Software (LI-COR).

### Immunoprecipitation and mass spectrometric analysis of intracellular tau

Tau was immunoprecipitated from 1 mg of total protein extracted from iPSC-derived neurons (120 DIV) using a polyclonal anti-tau antibody (Dako Cytomation; A0024). Immunoprecipitated samples were analysed by western blot using a monoclonal tau antibody (MN1000; ThermoFisher Scientific) or stained with colloidal blue (Invitrogen; LC6025). Bands that corresponded to tau by western blot analysis were excised from the colloidal blue SDS-PAGE. Peptide masses of digested protein samples were determined using a Bruker ultrafleXtreme Maldi mass spectrometer in reflectron mode and ms/ms fragmentation performed in LIFT mode. Data analysis was with FlexAnalysis, BioTools and ProteinScape software (Bruker). Database searches of the combined mass fingerprint-ms/ms data were performed using Mascot (http://www.matrixscience.com).

### Confocal and 3D-STED microscopy and image analysis

Standard confocal images of fixed and immunostained cells were acquired with an Olympus Inverted FV1000 confocal microscope and processed using Fiji software (Schindelin et al., 2012). 3D-STED imaging was performed on a custom built, dual color, beam scanning system with gated detection optically identical to the instrument described in (Bottanelli et al., 2016). For image analysis of colocalisation of tau and MAP2, Pearson’s R correlation was calculated using the Coloc2 plugin for Fiji software (https://imagej.net/Coloc2). To quantify nuclear invaginations, at least 5 imaging fields from three independent experiments for genotype were analysed using a custom plugin for the Fiji. See Fig S3 for details.

### Live imaging of microtubule dynamics

Neurons were grown to 120 DIV in individual μ-Dish 35 mm dishes (Ibidi; 80136) and transfected with a plasmid encoding for GFP-EB3 (Addgene plasmid # 56474). 48 hours after transfection, neurons were imaged live using a Leica SP5 microscope equipped with a controlled environment chamber (37 C; 5% CO_2_). Images were acquired at resonant scanning with a 63x objective (1frame/sec). Movies were analysed using the plusTipTracker software (Applegate et al., 2011).

### Analysis of human post-mortem cerebral cortex

Human brain sections were obtained from the Queen’s Square Brain Bank, Institute of Neurology, University College London. Control brains included one male (age 71) and one female (age 56). FTD-*MAPT* IVS10+16 brains were from two males (age 52 and 66). Nuclear lamina invaginations were quantified after 3,3′-Diaminobenzidine (DAB) staining of LaminB1. Nuclei were scored from 20 randomly acquired imaging fields, from each post-mortem sample, by three observers blinded to sample identity. The percentage of nuclei with invaginations relative to total nuclei was calculated for each imaging field (see **Fig. S4C**). For fluorescent immunostaining of laminB1 and tau, following incubation with secondary antibodies and DAPI nuclear counterstain, images were acquired using an Olympus Inverted FV1000 confocal microscope.

### Nucleocytoplasmic transport assay

Nucleocytoplasmic trafficking was analysed by infection of 120 DIV human iPSC-derived neurons with the pLVX-EF1alpha-2xGFP:NES-IRES-2xRFP:NLS construct (Addgene #71396; Mertens et al., 2015). After 6 days, neurons were fixed and immunostained for β3-tubulin and GFP. Only cells positive for neuron-specific β3-tubulin were considered. The nuclear to cytoplasmic ratios of both GFP and RFP (nucRFP:cytRFP and nucGFP:cytGFP) were calculated separately using the integrated density of ROIs drawn within and outside the nucleus (see **Fig.S5** for details).

## ACKNOWLEDGEMENTS

FJL’s group is supported by a Wellcome Trust Senior Investigator Award, the Alborada Trust’s funding of the Alzheimer’s Research UK Stem Cell Research Centre and the IMI program StemBANCC. SPJ is a Wellcome Trust Senior Investigator. SW and JH received funding from the National Institute for Health Research University College London Hospitals Biomedical Research Centre and SW by an ARUK Senior Research Fellowship (ARUK-SRF2016B-2). Research in SPJ and FJL’s groups benefits from core support to the Gurdon Institute from the Wellcome Trust and Cancer Research UK. The research leading to these results has received support from the Innovative Medicines Initiative Joint Undertaking under grant agreement n° 115439, resources of which are composed of financial contribution from the European Union’s Seventh Framework Programme (FP7/2007-2013) and EFPIA companies’ in-kind contribution. This report reflects only the author’s views and neither the IMI JU nor EFPIA nor the European Commission are liable for any use that may be made of the information contained therein.

